# Orthohantavirus infection in two rodent species that inhabit wetlands in the central-east region of Argentina

**DOI:** 10.1101/2022.01.05.475175

**Authors:** Malena Maroli, Carla M. Bellomo, Rocío M. Coelho, Valeria P. Martinez, Carlos I. Piña, Isabel E. Gómez Villafañe

**Affiliations:** Centro de Investigación Científica y de Transferencia Tecnológica a la Producción— Consejo Nacional de Investigaciones Científicas y Técnicas. Facultad de Ciencia y Tecnología, Universidad Autónoma de Entre Ríos, Diamante (3105), Entre Ríos, Argentina; Instituto Nacional de Enfermedades Infecciosas Administración Nacional de Laboratorios e Institutos de Salud Dr. Carlos G. Malbrán, Buenos Aires, Argentina; Instituto de Ecología, Genética y Evolución de Buenos Aires (CONICET-UBA). Facultad de Ciencias Exactas y Naturales. Universidad de Buenos Aires. Ciudad Autónoma de Buenos Aires, Argentina

**Keywords:** Emerging diseases, HPS, *Oligoryzomys*, *Oxymycterus*, Sigmodontine, Zoonoses

## Abstract

Hantavirus pulmonary syndrome (HPS) is an emerging infectious disease caused by orthohantaviruses associated to rodents of the Cricetidae family, Sigmodontinae subfamily, in the American continent. Previous research carried out in central-east region of Argentina, recorded potential orthohantavirus host rodents in diverse environments, but infected rodents were particularly present on Paraná wetlands islands. The aims of this research were (1) to determine the orthohantavirus host in the rodent community focused on islands of Paraná River Delta, an endemic zone of HPS, (2) to identify temporal and spatial factors associated with orthohantavirus prevalence variations, (3) to compare the individual characteristics of seropositive and seronegative rodents and, (4) to explore the association between orthohantavirus seroprevalence and rodent community characteristics in the Paraná River Delta, central-east region of Argentina. Capture of small rodents was done between August 2014 and May 2018 on seven islands located in central-east region of Argentina. In this HPS endemic zone, 14.9% of *Oligoryzomys flavescens* and 1.5% of *Oxymycterus rufus* of the sampled rodents had antibodies against orthohantavirus. The individuals that were more likely to become seropositive were the reproductively active adult males. Even though *O. flavescens* inhabit all islands, the seropositive individuals were only present in two of these, suggesting spatial heterogeneity in the viral distribution. We found that two months later of periods with low temperature, seroprevalence increased probably due to a higher proportion of adults in the population. Additionally, higher seroprevalence was associated with greater diversity of the rodent assemblage. This association could support the idea that a rescue effect or amplification of the prevalence of orthohantavirus would be taking place by means of secondary host as *O. rufus*, a novelty for this species and for the region. This finding may be significant if one takes into account that *O. rufus* was the second most abundant species in the area of islands studied and is one of the most abundant species on the islands and riparian sectors of the study zone. In conclusion, the relative risk of HPS could be high on wetlands of Paraná River Delta in the central-east region of Argentina where several favourable factors for the transmission of orthohantavirus are combined, such as the presence of several host species, two of them numerically dominant, high percentages of infection and a high degree of occupational exposure of the human population due to rural activities, the most frequently associated nationwide with HPS.

**Synopsis:** Hantavirus pulmonary syndrome (HPS) is an emerging infectious disease endemic of the American continent transmited by rodents. The aim of this research was to determine hosts species of orthohantaviruses in the rodent community on islands of Paraná River Delta, an HPS endemic zone of Argentina. We recorded the 14.9% of *Oligoryzomys flavescens* and 1.5% of *Oxymycterus rufus* with antibodies against orthohantavirus, which were principally reproductively active adult males. Seroprevalence increased after periods of low temperatures, probably due to the mortality of juveniles and survival of adults in the population. Additionally, the highest percentage of seropositive rodents occurred in times with a greater diversity of the rodent assemblage. This association could support the idea of amplification of the prevalence of orthohantavirus would be taking place by means of *O. rufus* infected, a novelty for this species and for the region. In conclusion, HPS risk could be high on wetlands of Paraná River Delta in the central-east region of Argentina where several favourable factors for the transmission of orthohantavirus are combined, such as the presence of several host species, high percentages of infection and a high degree of occupational exposure of the population due to rural activities.

## Introduction

Hantavirus pulmonary syndrome (HPS) is an emerging infectious diseases caused by orthohantavirus associated to rodents of Cricetidae family, Sigmodontinae subfamily, in The Americans [1–3]. The highest proportion of cases occurred in the Southern Cone of south America [4]. Orthohantaviruses are transmitted among rodents through aggressive encounters and by inhalation of aerosols contaminated with the virus [5,6], therefore, the transmission depends on the abundance, social structure of the community and behaviour of the species and individuals [7,8]. Ocasionally, spill-over events to non-traditional host can result in dead-end host, host-jumps or host-switch occurrences [9].

Ostfeld and Keesing [10] proposed the dilution effect hypothesis, originally to Lyme diseases, with the idea that the increase of the species diversity produce a decrease of the prevalence in the reservoir. Later, this hypothesis began to debated to Pfäffle et al. [11] in other host-pathogen systems, including orthohantavirus. On the contrary, the rescue effect indicates that if many species in the community tend to be highly competent reservoirs of a pathogen, when the specific diversity is high, the prevalence of the disease may increase [12,13]. Other researches have an intermediate position, indicating that species diversity can act in various ways on the determinants of transmission and can cause both increases and decreases in transmission [14–16] being a local phenomenon that depends on the specific composition of the reservoir, host, and its ecology, rather than on the patterns of species diversity [17].

In Argentina, four HPS endemic regions were identified; being Delta and Paraná islands ecoregion one of the most affected settings of the Central region (region with 36.7% of total country cases), particularly in rural and natural environmental, where around 23.9% cases occurs with high case-fatality rate values [18, 19]. Additionally, previous research carried out in diverse environmental settings inside this ecoregion recorded potential hantavirus host rodents in grassland, continental forest, riparian forest and islands [20, 21], but infected rodents were particularly present on the island [21, 22].

On this way, the aims of this research were (1) to determine the orthohantavirus host in the rodent community focused on islands of Paraná River Delta, an endemic HPS zone; (2) to compare the individual characteristics of rodents with and without antibodies against orthohantavirus; (3) to identify temporal and spatial factors associated with orthohantavirus prevalence variations, and (4) to explore the association between orthohantavirus circulation and rodent community characteristics.

## Methods

### Study area

The study was conducted on seven islands located inside a mosaic of wetlands, in the Paraná River Delta, central-east region of Argentina (Entre Ríos and Santa Fe provinces, Fig. 1). Two of the islands belong to the Pre-Delta National Park (PDNP, 32° 03’ 43”S; 60° 38’ 39”W), one of Islas de Santa Fe National Park (ISFNP, 32°16’44, 87’’ S, 60°43’12, 00’’ W) and the others four are private islands with livestock activity.

**Figure 1.**
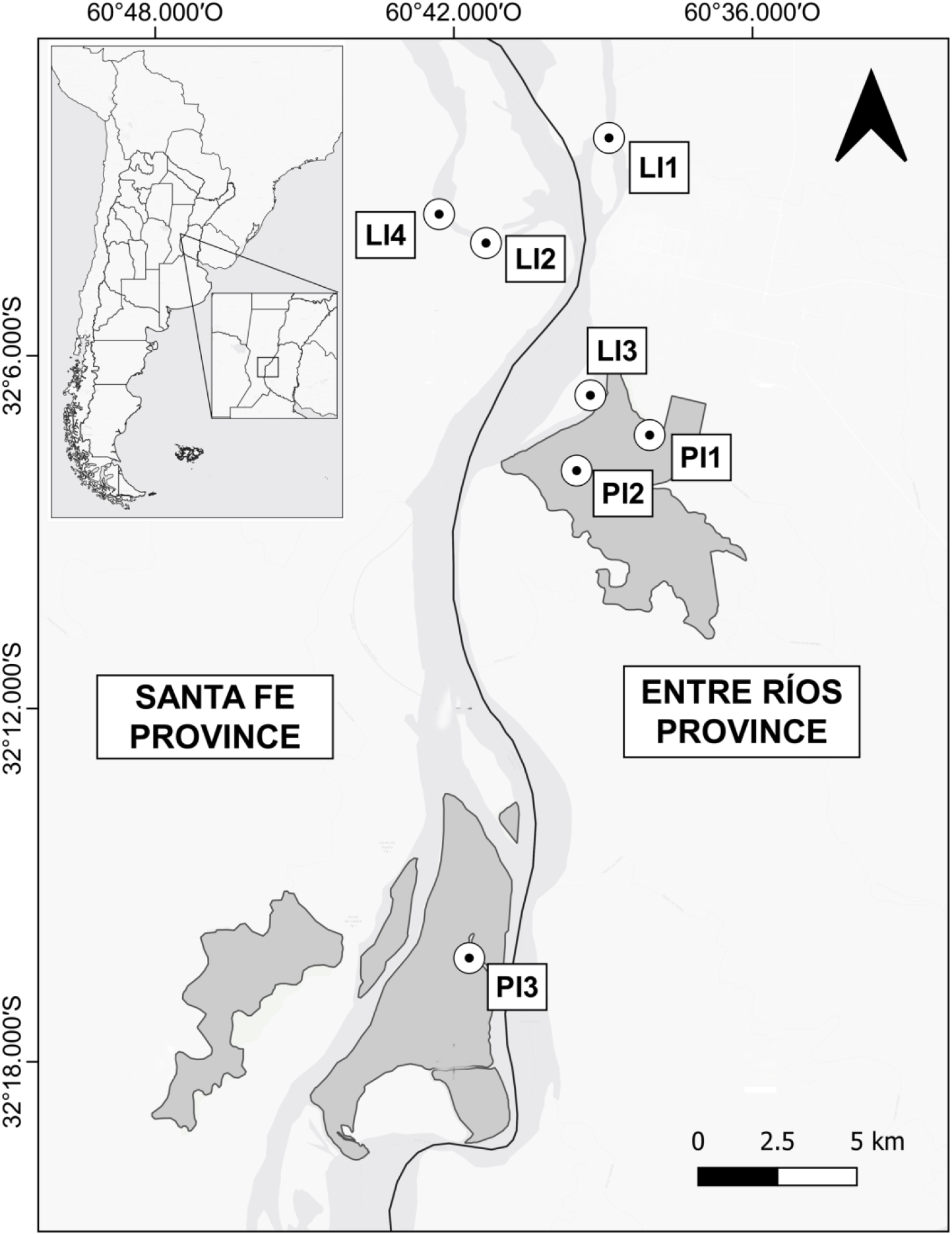
Location of the seven islands studied in Paraná River Delta, central-east region of Argentina (Entre Ríos and Santa Fe provinces; black solid line: limit between provinces and navigation channel of Paraná River). Two islands belong to Pre-Delta National Park (PI1, PI2), one belong to Islas de Santa Fe National Park (PI3) and four private islands with livestock activity (LI1, LI2, LI3, LI4). Islands PI1 and PI3 presented seropositive individuals.

### Rodent Surveys

Captures of small rodents were done quarterly between August 2014 and May 2018. Between 25 and 50 Sherman live traps (23 × 8 × 9.5 cm) baited with a mixture of peanut butter, fat and rolled oats were placed at 10m intervals in the levee of each island for three consecutive nights. The species, according to external characteristics [23], sex, total length, weight and reproductive status [24] of the captured individuals were recorded. Each individual was marked with an individually numbered ear tag (National Band and Tag Co., USA). Individuals were determined as adults based on their total lengths [23, 25]. For each trapping session and island, we calculated trap success (TS) as: number of individuals (not counting recaptures) /100 traps nights (TN). Also we account community characteristics as species richness (as number of different species), Shannon-Wiener diversity index following Magurran et al. [26] and relative abundance of host rodents (rodent species recorded with antibodies against orthohantavirus in this study).

### Environmental factors

Seventeen weather variables and fourteen hydrological variables associated with the temperature, rainfall, Oceanic Niño Index, river water level, number of days since last flooding event, frequency of flood and months elapsed since the evacuation level alert were recorded throughout the study period (see details in [20]).

### Orthohantavirus

Rodent blood samples were serologically screened by IgG ELISA using the N-ANDES hantavirus recombinant nucleocapsid protein [27]. IgG antibodies were detected in rodent sera diluted 1:400 using peroxidase-labeled affinity-purified IgG antibodies anti *Peromyscus leucopus* and anti-rat conjugate for *Rattus rattus* in conjunction with the ABTS Microwell Peroxidase Substrate System. The adjusted optical density for a reaction was the optical density of the well coated with the test antigen less the optical density of the well coated with the comparison antigen. We used antibody presence (seropositive individual) as an indication of orthohantavirus circulation in the population, however, it was not possible to infer whether the rodent was a reservoir or the antibodies were generated in response to a spillover. We calculated the seroprevalence as the percentage of seropositive individuals of the total analysing.

Animals were handled according to Argentinian National Law number 14.346 for the protection of animal welfare (Penal Code) and followed international guidelines appropriate for handling orthohantavirus reservoirs [28,29].

### Statistical analysis

Association among presence/absence of antibodies against orthohantavirus in *O. flavescens* individuals (binary response variable) and total length, body mass and sex (explanatory variables) was analyzed by a Bernoulli Generalized Linear Model (GLM), with cloglog link function [30, 31] and Laplace approximation method [30, 31]. The same analysis was repeated with only *O. flavescens* males including total length, body mass and reproductive condition (active, non-active) as explanatory variables. To assess the accuracy of the selected model we applied the kappa index [33] using *PresenceAbsence* package of R [34].

Association among orthohantavirus seroprevalence in *O. flavescens* and environmental factors and community characteristics were analyzed by means of GLM with binomial family distribution of errors and logit link [29].

For all models we assessed the association between all predictor variables using the Variance Inflation Factors (VIF>5) of *car* package [35,36]. Models were based on Akaike’s information criterion corrected for small sample size [37] with *AICcmodavg* [38] and *MuMIn* [39] packages of R. All statistical analyses were performed with R version 3.5.1 [40].

## Results

Orthohantavirus seroprevalence was 14.9% in *Oligoryzomys flavescens* (17/114) and 1.5% for *Oxymycterus rufus* (2/130). These two rodent species were the most abundant (47.1/100 TN, 164 *O. flavescens* and 34.8/100 TN, 130 *O. rufus*) followed for *Akodon azarae* (15.5/100 TN, 54 individuals), *Holochilus chacarius* (1.1/100 TN, 4 individuals), *Calomys callidus* (0.9/100 TN, 3 individuals), *Calomys laucha* (0.3/100 TN, 1 individual) and *Oligoryzomys nigripes* (0.3/100 TN, 1 individual) which did not present antibodies against orthohantavirus.

*O. flavescens* were captured in the seven islands at least once; however, seropositive individuals were captured only in two of the three islands of the national parks (PI1 and PI3 of the Fig. 1, Table 1). On PI1 rodents were captured in 10 of 13 samplings which were perform along the period (since winter 2017 to summer 2018), and seropositive individuals were detected in four of these 10 samplings in which seroprevalence ranged from 10% to 50% (Table 1). In the other island (PI3), the *O. flavescens* individuals were captured in two of six samplings (autumn and winter) and seropositive individuals were presented in autumn 2017, with a seroprevalence of 10% (Table 1).

**Table 1.** Details of seasons and islands where *Oligoryzomys flavescens* and *Oxymycterus* rufus were detected with antibodies against orthohantavirus.

| Trap seasons | Island | % *Oligoryzomys flavescens* |  | % *Oxymycterus rufus* |  |
| --- | --- | --- | --- | --- | --- |
|  |  | Seropositive | Tested | Seropositive | Tested |
| Winter 2017 | IP1 | 18.75 | 16 |  |  |
| Spring 2017 | IP1 | 30.00 | 10 |  |  |
| Summer 2017 | IP1 | 26.31 | 19 | 3.57 | 28 |
| Summer 2018 | IP1 | 50.00 | 2 | 22.22 | 9 |
| Autumn 2017 | IP3 | 10.00 | 50 |  |  |

*O. rufus* were captured in three of the seven studied islands (PI1, PI2 and LI3 of Fig. 1) but the two seropositive individuals were recorded on the same island (PI1 of the Fig. 1) and coincided with the island of seropositive *O. flavescens* (Table 1). One of them was first captured in October 2017, and at that time, it was seronegative. After two months (December 2017) this individual was recaptured and it could be verified that it had already acquired a reproductive condition and that it had seroconverted. Three months later (March 2018) it was recaptured for the third time, showing that it remained being seropositive. The other seropositive *O. rufus* was an adult male (non reproductive) captured on March 2018.

At the individual level for *O. flavescens*, seropositivity was higher in *O. flavescens* males (15.6%) than females (1.5%; Table 2 and 3), and the probability to find seropositive individuals was higher in longer individuals (total mean body length of seropositive individuals= 216.8 ± 2.7 mm; total mean body length of seronegative individuals= 193.8 ±1.6 mm; Table 2 and 3). The model has a moderate degree of concordance between observed and predicted values (Kappa index of 0.51± 0.12). The only seropositive *O. flavescens* female was a not reproductive adult (194 mm and 21 g). This female was captured on IP1 (Fig. 1) island together with other infected males in the summer of 2017. When the GLM statistical analysis was focused only on *O. flavescens* males the probability of orthohantavirus seroprevalence was greater in reproductive active (38.23%) than non-reproductive (3.22%) individuals, and in longer individuals (total mean body length of seropositive males= 212.2 ± 1.98; total mean body length of seronegative males= 189.1± 0.22 mm; Table 2 and 3). The model has a moderate degree of concordance between observed and predicted values (Kappa index of 0.52 ± 0.10).

**Table 2.**
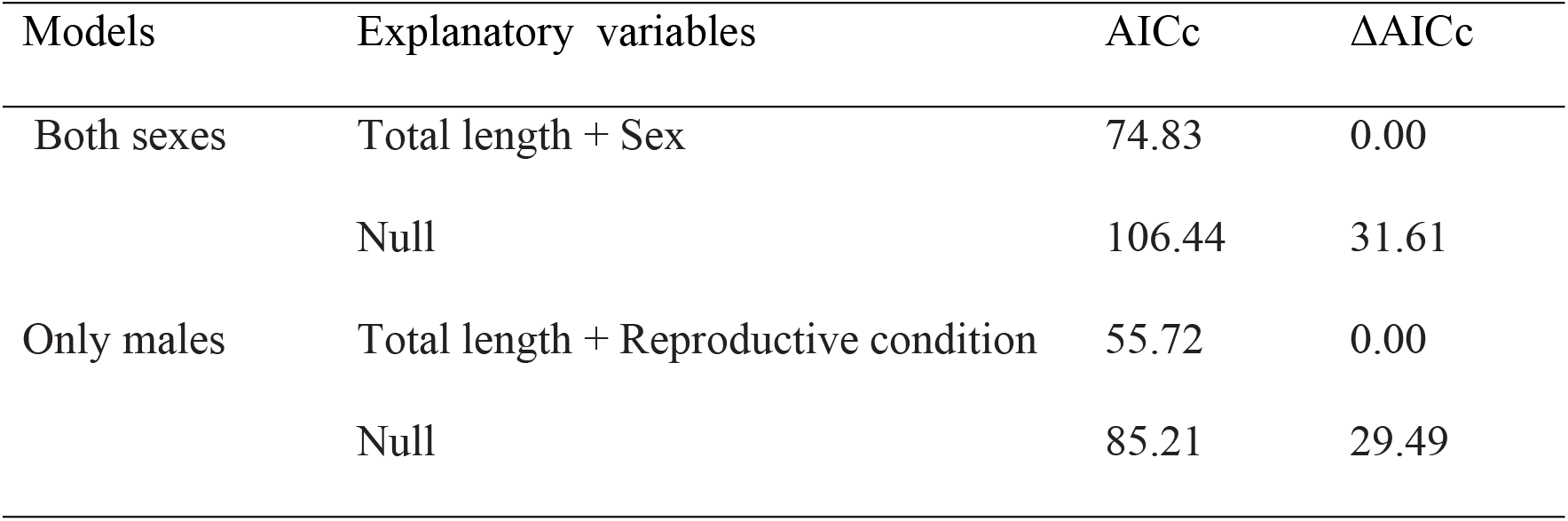
Selected models to explain the seropositive *Oligoryzomys flavescens* in relation to individual characteristics analysing both sexes together and only males, on Paraná River Delta islands, Argentina. AICc: Akaike’s criterion corrected for small samples; ΔAICc: difference of the AICc value of each model from the AICc value of the best model.

| Models | Explanatory variables | AICc | ΔAICc |
| --- | --- | --- | --- |
| Both sexes | Total length + Sex | 74.83 | 0.00 |
|  | Null | 106.44 | 31.61 |
| Only males | Total length + Reproductive condition | 55.72 | 0.00 |
|  | Null | 85.21 | 29.49 |

**Table 3.**
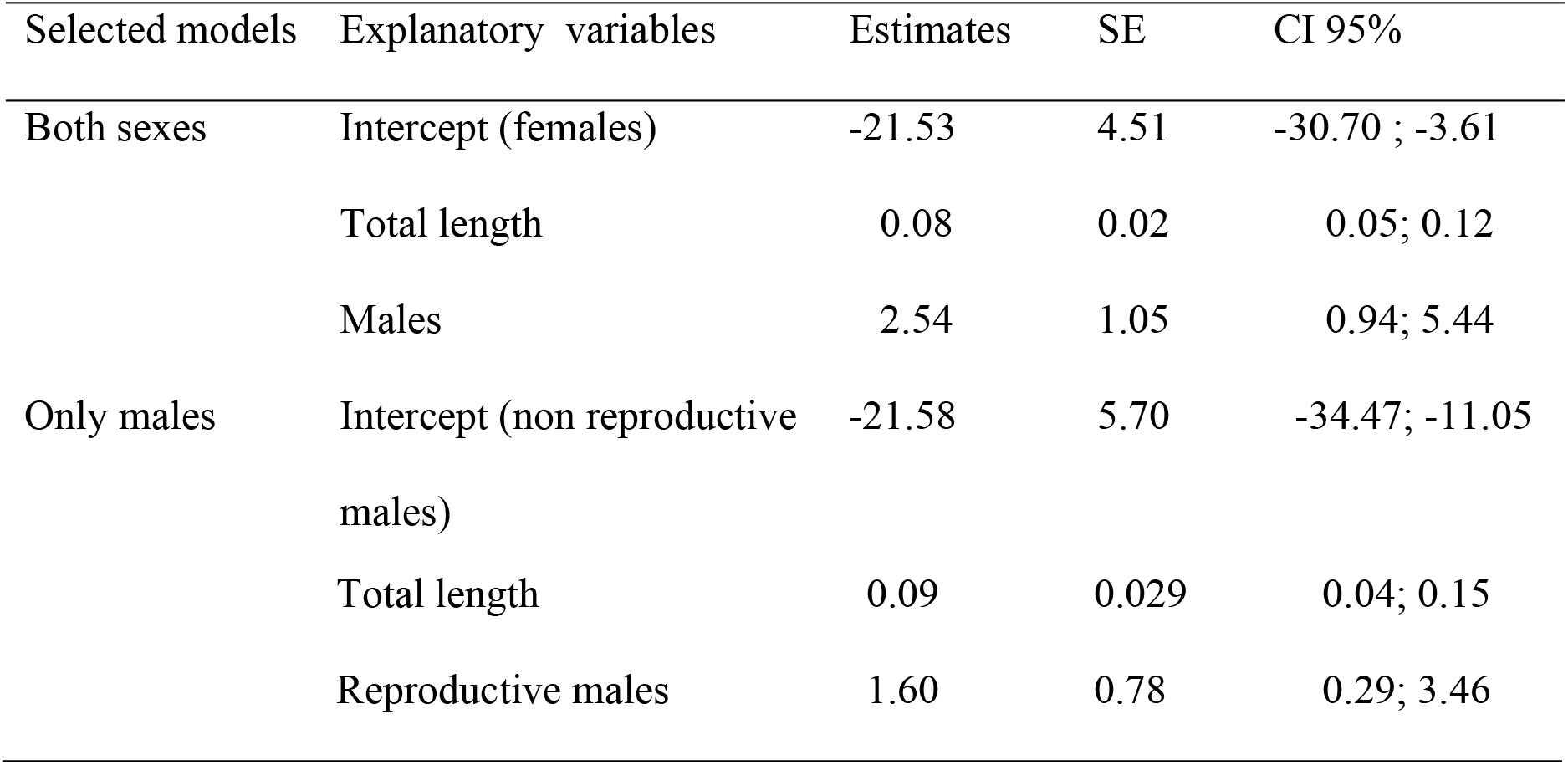
Estimates (log-Mean scale), standard error (SE) and 95% confidence interval limits (CI 95%) for selected models that explain *Oligoryzomys flavescens* seropositives in relation to individual characteristics, on Paraná River Delta islands, Argentina.

At population level, three models were selected that explained the 55.9%, 51.1% and 41% of variance, respectively (Table 4). The highest seroprevalence of orthohantavirus in *O. flavescens* occurs when the lowest ONI values are present, extreme minimum monthly temperature 60 days previous to trapping, and in time interval and on certain islands with the highest specific diversity (Shannon Wiener Index, Table 5).

**Table 4.** Selected model to explaining variation in orthohantavirus seroprevalence in relation to environmental and community characteristics on Paraná River Delta islands, Argentina. AICc: Akaike’s criterion corrected for small samples; ΔAICc: difference of the AICc value of each model from the AICc value of the best model.

| Models | Explanatory variables | AICc | ΔAICc |
| --- | --- | --- | --- |
| Environmental model 1 | ONI | 22.31 | 0.00 |
| Environmental model 2 | Minimum monthly temperature 60 days previous to trapping | 23.28 | 0.98 |
| Environmental null model |  | 30.89 | 8.59 |
| Community model | Rodent diversity | 35.53 | 0.00 |
| Community null model |  | 45.27 | 9.74 |

**Table 5.** Estimates (log-Mean scale), standard error (SE) and 95% confidence interval limits (CI 95%) for selected models that explain variation in orthohantavirus seroprevalence in relation to environmental and community characteristics on Paraná River Delta islands, Argentina.

| Models | Explanatory variable | Estimate | SE | CI 95% |
| --- | --- | --- | --- | --- |
| Environmental model 1 | Intercept | -2.53 | 0.36 | -3.33; -<br>1.91 |
|  | ONI | -2.58 | 0.87 | -4.37; -<br>0.93 |
| Environmental model 2 | Intercept | -1.29 | 0.42 | -2.20; -<br>0.51 |
|  | Minimum monthly<br>temperature 60 days<br>previous to trapping | -0.30 | 0.09 | -0.49;-<br>0.12 |
| Community model | Intercept | -3.50 | 0.57 | -4.75; -<br>2.50 |
|  | Diversity | 3.33 | 1.034 | 1.41; 5.50 |

## Discussion

In this research, we confirmed that orthohantaviruses are circulating in rodent populations in the study area, mainly in the species *O. flavescens* and *O. rufus* that inhabit on islands of Central Argentina. Previous studies in this HPS endemic zone, identified two rodent species as host of pathogenic orthohantaviruses: *O. flavescens*, that carried the genotypes Lechiguanas, Buenos Aires and Plata genotypes in Argentina and Uruguay [21,41,42] and *O. nigripes* with the genotype lechiguanas [21].

Individuals of the species *O. flavescens* that were more likely to become infected were reproductively active adult males, in accordance with previous studies [21,44–48]. The global orthohantavirus seroprevalence recorded in *O. flavescens* had similar values to those presented by other researches (3.6% to 8% [49, 50, 51, 21]). However, specific seroprevalence detected in particular times and islands were very high, showing the possibility that populations of *O. flavescens* can reach high degrees of infection in the Delta of the Paraná River. Even though *O. flavescens* inhabit all islands, seropositive individuals were only found in three out of seven, suggesting a spatial heterogeneity in the virus distribution. This is consistent with other orthohantavirus-host systems that evidence the existence of spatial focality at different various scales [21,52–56].

Additionally, the orthohantavirus seroprevalence in the *O. flavescens* population showed temporal variation, detecting a maximum in mid-2017 (winter). This fact could be due to the mortality of juvenile rodents under adverse weather. Therefore, the proportion of adults increase, and, consequently also the proportion of infected individuals (inverse effect of the dilution effect through the presence of juveniles that are less infected than adults). In turn, the seroprevalence in this study was higher in periods of occurrence of ENSO-La Niña (which in south eastern South America causes drought conditions [57] but this aspect, opposite to the positive association between prevalence and precipitation found in other research [58] requires further study.

Analyzing the rodent community, seroprevalence of orthohantavirus in this islands system did not show any association with the abundance or the relative abundance of the reservoir *O. flavescens*. However, higher prevalence was detected associated with greater diversity of the rodent assemblage. This increase would support the idea that a rescue effect or amplification of the prevalence of hantavirus would be taking place in the rodent assemblage of the Paraná River Delta islands. The mechanism through which this amplification effect could act would be the presence of some species of rodents that until now were not considered a reservoir species (at least of the genotype in question). This species could begin to behave as competent hosts for the virus and, therefore, detect accidental infection in individuals; a phenomenon that in the long term could lead to a change of host for the virus. The fact that could reinforce the evidence of an amplification effect of the prevalence of hantavirus infection in the Paraná River Delta is the finding of seropositive individuals of *O. rufus* in the same island and moments that seropositives *O. flavescens* were recorded, and the presence of the species *O. nigripes. O. nigripes* was found carrying Lechiguanas virus in the same region [21] and it was described as the reservoir host of Juquitiba virus in previous works [62] De Olivera Renata et al 2014). In addition, the seropositive *O. rufus* that was recaptured twice after the first time, evidencing a seroconversion process between the first and the second sample, status that was confirmed in the third capture event, three months later. Although among the characteristics to define a host as a reservoir is the presence of the viral genome, other characteristics remain to be evaluated to define *O. rufus* as a reservoir: infecting viral variant, intraspecific transmissibility -assessed by detecting a high number of seropositive individuals- and absence of symptoms or other characteristics that could influence the fitness of infected individuals. It would also be important to determine whether this species is capable of infecting humans. This may be significant taking into account that *O. rufus* was the second most abundant species in this area of islands and is one of the most abundant species on other islands and riparian sectors of the Paraná River Delta [20,59].

Based on this result, and that other genetically related species infected with pathogenic variants have been detected, such as the species *Oxymycterus nasutus* in Uruguay (Araquara genotype [60]) and *Oxymycterus judex* in Brazil (Araquara genotype [61]), demonstrates the need to monitor this species to elucidate its role in the hantavirus-host system of the Paraná River Delta and evaluate potential risks for humans. The species *O. nigripes*, captured during this study, as details above, is a known reservoir of the Juquitiba pathogenic genotype in Brazil and northeastern Argentina [41,62], and was also described as a host of the pathogenic Lechiguanas virus in Central Argentina [21] however, it was not possible to monitor this species due to its very low abundance in the area.

In conclusion, several favourable factors for the transmission of orthohantavirus are combined, as the presence of several host species, two of them numerically dominant, high percentages of infection on some islands added to a high degree of human occupational and recreational activities in wild areas such as is known in these types of rural areas [18, 19, 62–64], which would mean that wetlands of Paraná River Delta in the central-east region of Argentina is a risk zone for humans to become infected, and therefore need active surveillance.

## Acknowledgements

We are thankful to CONICET and Universidad de Buenos Aires for financial support for this study; to Administración de Parques Nacionales for their research authorizations to work in Pre-Delta and Islas de Santa Fe National Parks and their logistic support; to Entre Ríos and Santa Fe provinces for extending research authorizations. We also thank to Eliana Burgos and Belén Crosignani for providing their help during the fieldwork.

## Author Contributions

IGV conceived and designed the research. MM performed the field work. CB, RC and VM performed the laboratory work. MM and IGV analyzed the statistical data. MM and IGV wrote the manuscript. All authors participated in the final version of the manuscript.

